# The Hepcidin Regulator Erythroferrone is a New Member of the Erythropoiesis–Iron–Bone Circuitry

**DOI:** 10.1101/2021.04.09.439174

**Authors:** Melanie Castro-Mollo, Sakshi Gera, Marc Ruiz Martinez, Maria Feola, Anisa Gumerova, Marina Planoutene, Cara Clementelli, Veena Sangkhae, Carla Casu, Se-Min Kim, Vaughn Ostland, Huiling Han, Elizabeta Nemeth, Robert Fleming, Stefano Rivella, Daria Lizneva, Tony Yuen, Mone Zaidi, Yelena Z. Ginzburg

**Affiliations:** Division of Hematology Oncology, Tisch Cancer Institute, Icahn School of Medicine at Mount Sinai, New York, NY; The Mount Sinai Bone Program, Departments of Medicine and Pharmacological Sciences, and Center for Translational Medicine and Pharmacology, Icahn School of Medicine at Mount Sinai, New York, NY; Center for Iron Disorders, University of California, Los Angeles (UCLA), Los Angeles, CA; Department of Pediatrics, Division of Hematology, and Penn Center for Musculoskeletal Disorders, Children’s Hospital of Philadelphia (CHOP), University of Pennsylvania, Perelman School of Medicine, Philadelphia, PA; Intrinsic Lifesciences, LLC, LaJolla, CA; Department of Pediatrics, Saint Louis University School of Medicine, St Louis, MO

**Keywords:** osteoporosis, bone formation, transgenic mice, thalassemia, osteoblasts

## Abstract

Erythroblast erythroferrone (ERFE) secretion inhibits hepcidin expression by sequestering several bone morphogenetic protein (BMP) family members to increase iron availability for erythropoiesis. We report that ERFE expression in osteoblasts is higher compared with erythroblasts, is independent of erythropoietin, and functional in suppressing hepatocyte hepcidin expression. *Erfe*^-/-^ mice display low–bone–mass arising from increased bone resorption despite a concomitant increase in bone formation. Consistently, *Erfe*^-/-^ osteoblasts exhibit enhanced mineralization, *Sost* and *Rankl* expression, and BMP–mediated signaling *ex vivo*. The ERFE effect on osteoclasts is mediated through increased osteoblastic RANKL and sclerostin expression, increasing osteoclastogenesis in *Erfe*^-/-^ mice. Importantly, *Erfe* loss in β–thalassemic (*Hbb*^*th3*/+^) mice, a disease model with increased ERFE expression, triggers profound osteoclastic bone resorption and bone loss. Together, ERFE exerts an osteoprotective effect by modulating BMP signaling in osteoblasts, decreasing RANKL production to limit osteoclastogenesis, and prevents excessive bone loss during expanded erythropoiesis in β–thalassemia.

## INTRODUCTION

It has become increasingly clear that both erythropoiesis and skeletal homeostasis are susceptible to changes in iron metabolism, especially during stress or ineffective erythropoiesis. Diseases of ineffective erythropoiesis, such as β–thalassemia, one of the most common forms of inherited anemia worldwide^1^, are thus associated with bone loss, primarily at cortical sites^2,3^. β–thalassemia results from β-globin gene mutations that cause ineffective erythropoiesis, splenomegaly, and anemia^4–7^. Patients with homozygous mutations have either red blood cell (RBC) transfusion-dependent β–thalassemia major (TDT) or a relatively milder anemia, namely non-transfusion-dependent β–thalassemia intermedia (NTDT).

Both TDT and NTDT generally present with anemia and iron overload, requiring iron chelation therapy. Surprisingly, however, TDT patients show more marked decrements in bone mineral density (BMD) compared with NTDT, despite chronic RBC transfusion that suppresses expanded and ineffective erythropoiesis. Optimization of RBC transfusion has reduced the frequency of overt bone disease, such as frontal bossing, maxillary hyperplasia and limb deformities, and importantly, has enabled prolonged survival^8^. Nonetheless, growth patterns have not significantly improved^9^, and low bone mass remains a frequent, significant, and poorly understood complication even in optimally–treated patients. As such, β–thalassemia-induced bone disease has warranted formal guidelines for management^10^.

Proposed mechanisms of bone loss in β-thalassemia include direct effects of abnormal erythroid proliferation^11,12^, increased circulating erythropoietin (Epo)^13^, iron toxicity^14^, oxidative stress^15^, inflammation^16^, and changes in bone marrow adiposity^17^. Strong negative correlations between BMD and systemic iron concentrations^18^ and the profound bone loss noted in patients with hereditary hemochromatosis^19^ underscore the premise that BMD and iron homeostasis may be associated causally. However, mice lacking the transferrin receptor, TFR2, display *increased* rather than decreased bone mass and mineralization despite iron overload^20^. These latter findings prompted us to take a fresh look at the mechanisms underpinning bone loss in diseases of iron dysregulation.

Erythroferrone (ERFE), a protein secreted by bone marrow erythroblasts, is a potent negative regulator of hepcidin^21^, which, in turn, inhibits iron absorption and recycling^22^. Hepcidin suppression enables an increase in iron availability during stress erythropoiesis. Very recently, ERFE has been shown to bind and sequester certain members of the bone morphogenetic protein (BMP) family, prominently BMP2, BMP6 and the BMP2/6 heterodimer^23,24^. BMPs stimulate bone formation by osteoblasts during skeletal development, modeling, and ongoing remodeling^25^. We thus hypothesized that, by modifying BMP availability, ERFE may be a key player in the newly discovered erythropoiesis–iron–bone circuitry. As a result, ERFE may also be an important link between altered iron metabolism, abnormal erythropoiesis, and bone loss in β–thalassemia.

The known mechanism of ERFE action—BMP sequestration—predicts that ERFE loss, by enhancing BMP availability, may stimulate osteoblastic bone formation^26^. Alternatively, recent literature shows that loss of BMP signaling increases bone mass through direct osteoclast inhibition and Wnt activation^27–32^, predicting that ERFE loss would lead to decreased bone mass. Here, we demonstrate that global deletion of *Erfe* in mice results in a low–bone–mass phenotype, which is phenocopied in β–thalassemic mice lacking ERFE (i.e., *Hbb*^*th3*/+^;*Erfe*^-/-^ mice). Despite the osteopenic phenotype, we found that ERFE loss stimulated mineralization in cell culture. The net loss of bone in *Erfe*^-/-^ mice in the face of a pro–osteoblastic action could therefore only be attributed to a parallel increase in bone resorption, which we found was the case in both *Erfe*^-/-^ and *Hbb*^*th3*/+^;*Erfe*^-/-^ mice. Furthermore, the increase in osteoclastogenesis was osteoblast–mediated exerted *via* an increased expression of *Sost* and *Tnfsf11*, the gene encoding RANKL. Together, our data provide compelling evidence that ERFE loss induces BMP–mediated osteoblast differentiation, but upregulates *Sost* and *Tnfsf11* to increase osteoclastogenesis with net bone loss. Therefore, high ERFE levels in β-thalassemia are osteoprotective and prevent the bone loss when erythropoiesis is expanded.

## METHODS

### Mouse Lines

C57BL/6 and β-thalassemic (*Hbb*^*th3*/+^) mice^33^ were originally purchased from Jackson Laboratories. *Erfe*^-/-^ mice were a generous gift from Tomas Ganz (UCLA)^21^. Progeny of *Erfe*^-/-^ mice crossed with *Hbb*^*th3*/+^ yielded *Hbb*^*th3*/+^;*Erfe*^-/-^ mice. The mice have been backcrossed onto a C57BL/6 background for more than 11 generations. All mice had *ad libitum* access to food and water and were bred and housed in the animal facility under AAALAC guidelines. Experimental protocols were approved by the Institutional Animal Care and Use Committee at Icahn School of Medicine at Mount Sinai.

### Skeletal Phenotyping

Skeletal phenotyping was conducted on 6–week–and/or 5–month–old male mice, unless otherwise noted. Mice were injected with calcein (15 mg/kg, Sigma C0875) and xylenol orange (90 mg/kg, Sigma 52097), at days −8 and−2, respectively, prior to sacrifice. Briefly, for histomorphometry, the left femur, both tibias, and L1-L3 were fixed in neutral buffered formalin (10%, v/v) for 48 hours at 4°C; transferred to sucrose (30%, w/v) at 4°C overnight; and embedded and sectioned at −25°C (5-6 μm thick sections, 10X)^34^. Unstained sections were analyzed by fluorescence microscopy (Leica Upright DM5500) to determine the mineralizing surface and interlabel distance using image J. Von Kossa staining of sections was used to quantify fractional bone volume (BV/TV) and trabecular thickness (Tb.Th). Tartrate-resistant acid phosphatase (ACP5) staining (Sigma 387A) was used to identify osteoclasts, counterstained with aniline blue using Olympus Stereoscope MVX10 (1X). Images were analyzed by TrapHisto and

OsteoidHisto^35^. On the day of sacrifice, BMD was also measured in intact mice^36^. Frozen bone sections were incubated for 4 min at room temperature in Alkaline Phosphatase Substrate Solution ImmPACT Vector Red (Vector Laboratories). After washing with buffer, the sections were counterstained with hematoxylin (Vector Laboratories) and mounted with VectaMount AQ Mounting Medium (Vector Laboratories). Sections were visualized using Olympus BH-2 Microscope and images obtained with OMAX A35180U3 Camera were analyzed by ImageJ.

### Isolation and Culture of Bone Marrow Cells

Erythroblasts were isolated from bone marrow and purified using CD45 beads, as previously^37^ with minor modifications. Briefly, mouse femur was flushed, single-cell suspensions incubated with anti-CD45 magnetic beads (Mylteni), and erythroid lineage-enriched cells that flowed through the column were collected. Erythroid-enriched cells were incubated with antimouse TER119-phycoerythrin Cy7 (PE-Cy7) (BioLegend) and CD44-allophycocyanin (APC) (Tonbo, Biosciences). Non-erythroid and necrotic cells were identified and excluded from analyses using anti-CD45 (BD Pharmigen), anti-CD11b, and anti-Gr1 (APC-Cy7) (Tonbo, Biosciences) antibodies. Erythroid precursors were selected by gating and analyzed using TER119, CD44, and forward scatter as previously described^37^. Samples were analyzed on either FACSCanto I or LSRFortessa flow cytometer (BD Biosciences). To determine levels of *Erfe* mRNA expression in Epo–stimulated conditions, erythroblasts were cultured in the presence or absence of 20 U/ml Epo for 15 hours as described^38^. For osteoblast cultures, fresh bone marrow cells were seeded in 12–well plates (0.6×10^6^ cells *per* well) under differentiating conditions [αMEM, 10% FBS, 1% penicillin/streptomycin, 1 M β-glycerol phosphate, and 0.5 M ascorbic acid] for 21 days to induce the formation of mature, mineralizing osteoblast colonies, Cfu-ob, as before^39^. For osteoclast cultures, bone marrow hematopoietic stem cells (non–adherent) from wild type and *Erfe*^-/-^ were seeded in 6–well plates (10^6^ cells *per* well) in the presence of αMEM, 10% FBS, 1% penicillin/streptomycin and M-CSF (25 ng/mL, PeproTech) for 48 hours, followed by the addition of RANK-L (50 ng/mL, PeproTech) for 5 days. In experiments testing Epo responsiveness, 20 U/ml of Epo was added for the duration of the differentiation process, for 21 and 5 days in osteoblast and osteoclast cultures, respectively.

### Primary Hepatocyte Culture

Hepatocytes were isolated by perfusion with collagenase and liver digestion, as described previously^40^. Briefly, 0.025% (w/v) collagenase type IV (Gibco) and 5 mM CaCl_2_ was added to Leffert perfusion buffer containing 10 mM HEPES, 3 mM KCl, 130 mM NaCl, 1 mM NaH_2_PO_4_.H_2_O, and 10 mM D-glucose (Sigma). Live cells were purified by Percoll (Sigma) and plated in 6-well plates (0.25×10^6^ cells *per* well) in William’s Medium E (Sigma) supplemented with antibiotics and 5% fetal bovine serum (FBS) for 2 hours to allow the hepatocytes to attach. Cells were starved overnight with William’s Medium E lacking FBS, and were then treated for 6 hours with conditioned or control media from wild type or *Erfe*^-/-^ osteoblast and osteoclast cultures (day 6 and day 5, respectively) in the presence of 50% (v/v) William’s Medium E and 5% FBS.

### Quantitative PCR

RNA was purified from osteoblasts, osteoclasts, erythroblasts, and hepatocytes using PureLink RNA (Sigma) and analyzed with SuperScript III Platinum SYBR Green One-Step (Invitrogen). As previously described^41,42^, ΔΔCT values were used to calculate fold increases relative to β-actin, α-tubulin, and RLP4. Primers are listed in Table I.

**Table I.**
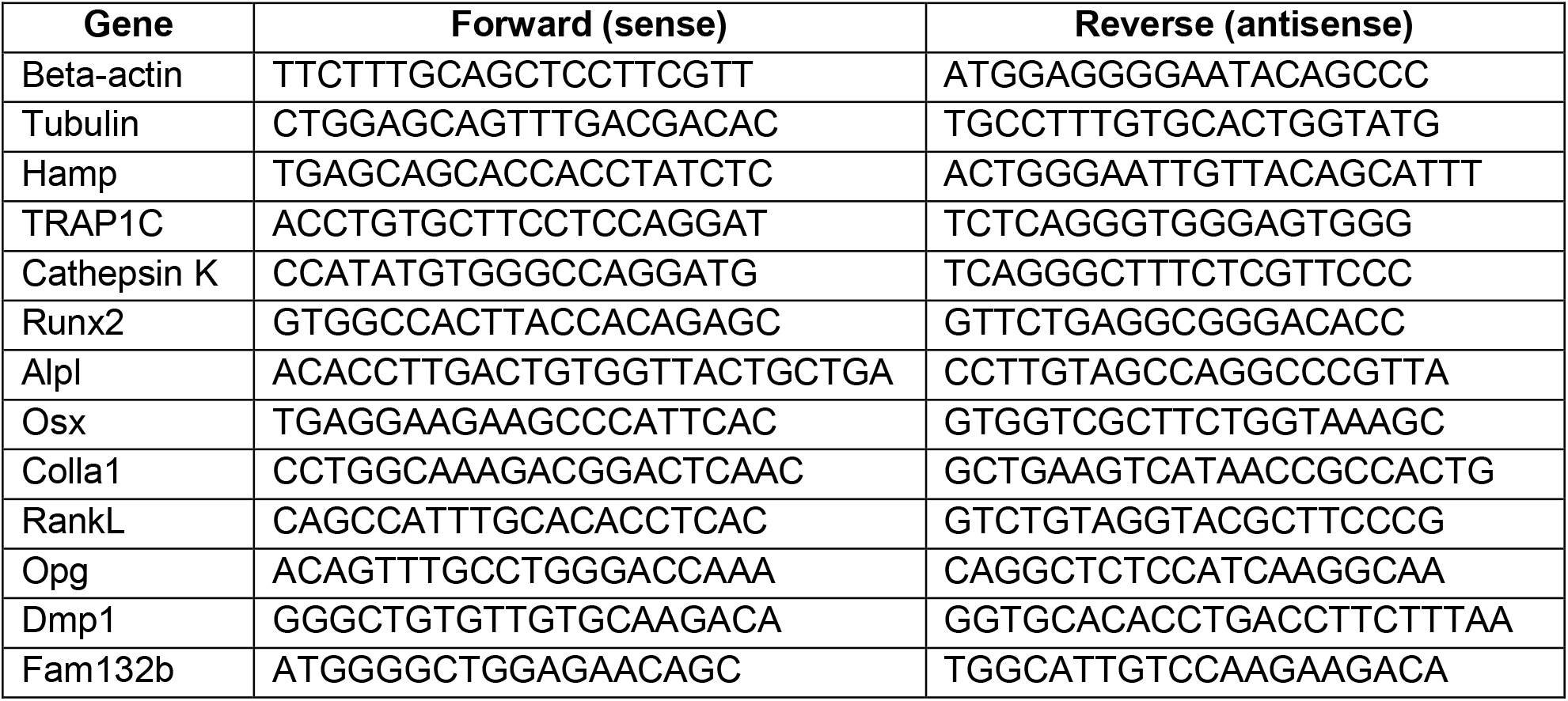

### Western Immunoblotting and ELISA

For Western immunoblotting, differentiated cells at day 3 were lysed in ice cold SDS page lysis buffer (2% SDS, 50 mM Tris-HCl, pH 7.4, 10 mM EDTA) with protease and phosphatase inhibitors. 20 μg of heat-denatured protein was loaded onto a 10% gel, run, and transferred onto a 0.4 μm nitrocellulose membrane (Thermo Scientific). After blocking with 5% BSA in Tris–buffered saline with 1% Tween-20 (TBS-T), the membranes were incubated with primary antibodies to signaling proteins (Table II) overnight at 4°C, washed, and incubated with the corresponding HRP–conjugated secondary antibodies at room temperature. Proteins were visualized using the ImageQuant LAS 4010 and quantified using Image J. Osteoblast supernatants from wild type and *Erfe*^-/-^ mice were collected and centrifuged for 10 min at 10,000 x g, and BMP2 (Abnova) and RANKL (R&D) concentrations were measured by ELISAs. Serum BMP2 concentration was determined using mouse BMP2 ELISA (abnova, KA0542), *per* manufacturers instructions. ERFE concentration in conditioned media was determined as described^43^ with the substitution of DELFIA europium–conjugated streptavidin for horseradish–peroxidase–conjugated streptavidin. Fluorescence was measured by CLARIOstar plate reader.

**Table II.**
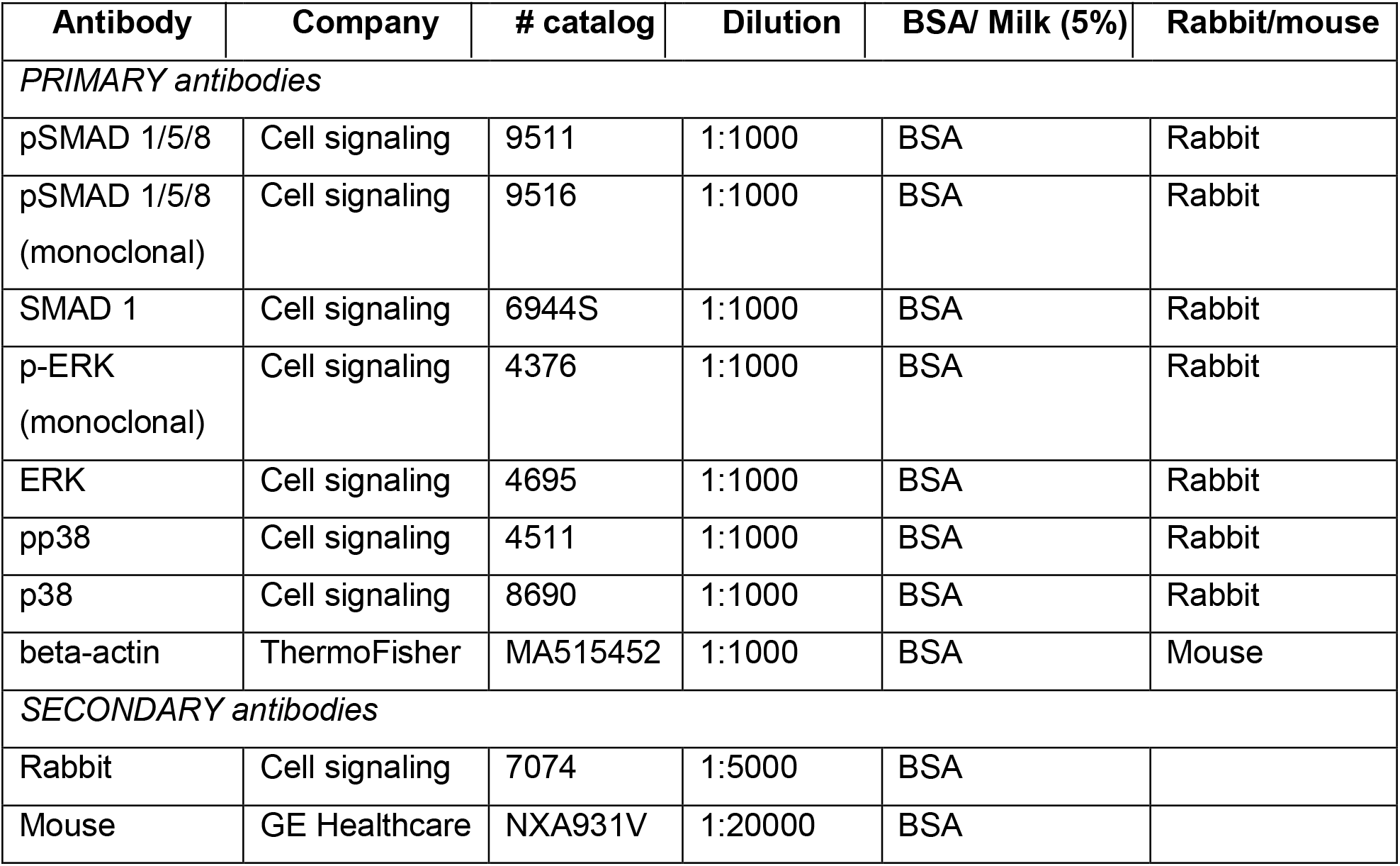

| Antibody | Company | # catalog | Dilution | BSA/ Milk (5%) | Rabbit/mouse |
| --- | --- | --- | --- | --- | --- |
| *PRIMARY antibodies* |  |  |  |  |  |
| pSMAD 1/5/8 | Cell signaling | 9511 | 1:1000 | BSA | Rabbit |
| pSMAD 1/5/8<br>(monoclonal) | Cell signaling | 9516 | 1:1000 | BSA | Rabbit |
| SMAD 1 | Cell signaling | 6944S | 1:1000 | BSA | Rabbit |
| p-ERK<br>(monoclonal) | Cell signaling | 4376 | 1:1000 | BSA | Rabbit |
| ERK | Cell signaling | 4695 | 1:1000 | BSA | Rabbit |
| pp38 | Cell signaling | 4511 | 1:1000 | BSA | Rabbit |
| p38 | Cell signaling | 8690 | 1:1000 | BSA | Rabbit |
| beta-actin | ThermoFisher | MA515452 | 1:1000 | BSA | Mouse |
| *SECONDARY antibodies* |  |  |  |  |  |
| Rabbit | Cell signaling | 7074 | 1:5000 | BSA |  |
| Mouse | GE Healthcare | NXA931V | 1:20000 | BSA |  |

### Complete Blood Counts

Peripheral blood (100 μL from each mouse) was collected from the retro-orbital vein in EDTA-coated tubes and analyzed by IDEXX Procyte Hematology Analyzer.

### Statistical Analyses

Data are reported as means ± SEM. Unpaired Student’s t-test was used to determine if differences between groups were significant at *P*<0.05.

## RESULTS

To understand if ERFE has a role in regulating skeletal integrity in health, we first studied the effect of ERFE loss on BMD and bone remodeling in adult *Erfe*^-/-^ mice, as well as in compound mutant mice in which the *Erfe* gene was deleted on a β-thalassemia *Hbb*^*th3*/+^ background.

Compared with wild type littermates, both 6–week–old and 5–month–old male *Erfe*^-/-^ mice showed significant reductions in whole body BMD, and BMD at mainly-cortical (femur and tibia) sites (Fig. 1A and 1B). However, in contrast to young mice, the older *Erfe*^-/-^ mice did not show a difference in lumbar spine BMD compared with wild type littermates. Interestingly, unlike hypogonadal bone loss, which is predominantly trabecular, the sustained reduction in femur and tibia BMD is consistent with prominent cortical loss seen in patients with β-thalassemia^2,3^.

**Figure 1:**
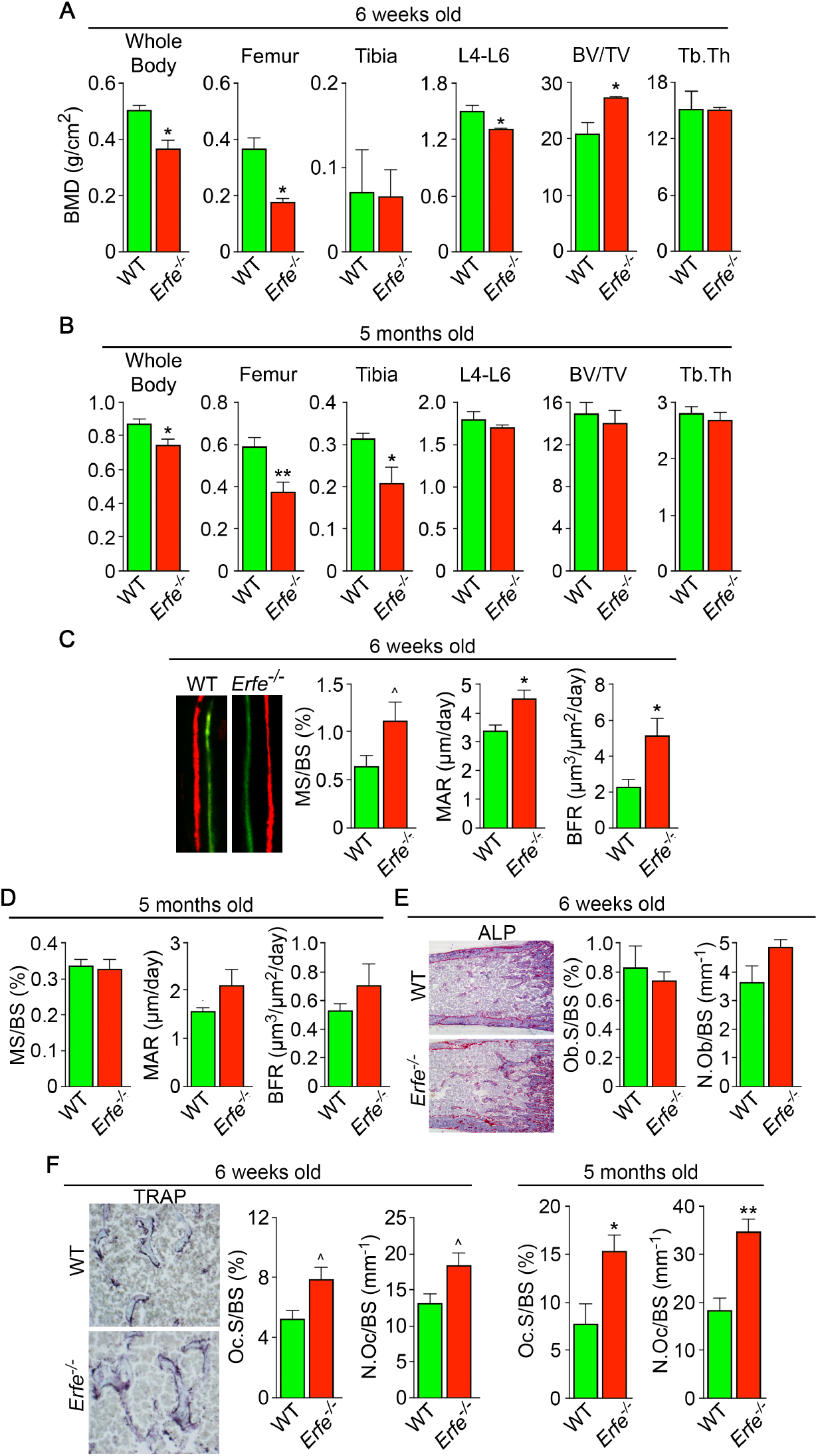
ERFE Loss Results in High Turnover Osteoporosis. Bone mineral density (BMD) measured in whole body, femur, tibia, and lumbar spine (L4-L6) along with bone volume (BV/TV) and trabecular thickness (Tb.Th) in growing (6–week–old) (**A**) and mature (5–month–old) (**B**) *Erfe*^-/-^ mice and wild type (WT) littermates. Dynamic histomorphometry following two i.p. injections of calcein (green) and xylenol orange (red) given at days −8 and −2, respectively. Representative dual labels from the epiphysis are shown, together with measured and derived parameters, namely mineralizing surface (MS) as a function of bone surface (BS), mineral apposition rate (MAR) and bone formation rate (BFR) 6–week–old (**C**) and 5–month–old mice (**D**). (**E**) Alkaline phosphatase staining (magenta) in sections of femura demonstrates no differences in osteoblast surface (Ob.S) and number (N.Ob) as a function of BS in 6–week–old *Erfe*^-/-^ relative to WT mice. (**F**) TRAP staining at the epiphysis showing both osteoclast surface (Oc.S) and number (N.Oc). Statistics: Mean ± SEM; unpaired two–tailed Student’s t–test; *P<0.05, **P<0.01; ^0.05<P<0.1, *N*=3–6 mice per group.

Bone resorption and bone formation are tightly coupled to maintain bone mass during each remodeling cycle^44^. Bone is lost when either both processes are increased—with resorption exceeding formation, as in hypogonadism—or when there is uncoupling in which formation decreases while resorption rises, as in glucocorticoid excess^44^. To differentiate between relative increases and uncoupling, we measured both formation and resorption in intact bone. Dynamic histomorphometry performed after the sequential injections of calcein and xylenol orange, which yielded dual fluorescent labels, allowed us to derive parameters of bone formation. We observed that mineralizing surface (MS), mineral apposition rate (MAR) and bone formation rates (BFR) were all increased in young *Erfe*^-/-^ mice, consistent with the pro–osteoblastic (anabolic) action of ERFE deficiency (see below) (Fig. 1C and 1D). No differences in MS, MAR, and BFR were noted in 5–month–old mice (Fig. 1D). We also analyzed alkaline phosphatase stained sections of femurs to find no difference in osteoblast surfaces (Ob.S) or osteoblast number (N.Ob) *per* bone surface (BS) in 5–week–old *Erfe*^-/-^ relative to wild type littermates (Fig. 1E).

Finally, to study whether an increase in osteoclastic bone resorption caused the notable reduction in BMD in *Erfe*^-/-^ mice, we measured TRAP–positive osteoclast surfaces (Oc.S) and number (N.Oc) *per* bone surface (BS). Both Oc.S/BS and N.Oc/BS were increased significantly in *Erfe*^-/-^ compared with wild type bones in older mice, and to a lesser extent, in younger mice (Fig. 1F). Thus, the overall low–bone–mass phenotype in *Erfe*^-/-^ mice primarily resulted from a relative increase in osteoclastic bone resorption over osteoblastic bone formation, suggesting that ERFE has a function in preventing skeletal loss. To confirm that decreased BMD in *Erfe*^-/-^ mice did not result from changes in erythropoiesis, we measured circulating red blood cells (RBCs) and reticulocytes, and bone marrow erythroblasts. We also measured spleen weight given the ubiquity of compensatory erythropoiesis that results in splenomegaly. Our results show no differences between 6–week–old wild type and *Erfe*^-/-^ mice (Supplementary Fig. 1), consistent with what has been previously reported in *Erfe*^-/-^ mice^21^.

To probe the mechanism of action of ERFE on osteoblastic bone formation and osteoclastic bone resorption, we first asked which cells in bone marrow produce ERFE, and whether secreted ERFE was functional. Intriguingly, time course studies in differentiating osteoblasts revealed that *Erfe* expression was 10– and 2–fold higher at 3 and 21 days of culture, respectively, compared with cultured erythroblasts--the only previously known source of ERFE in bone marrow (Fig. 2A). Furthermore, *Erfe* expression in mature osteoclasts was similar to cultured erythroblasts, with little expression in immature osteoclasts (Fig. 2B). Likewise, conditioned media from osteoblast cultures revealed increased ERFE concentration at 3 days with no differences in conditioned media from osteoclast cultures (Fig. 2C).

**Figure 2:**
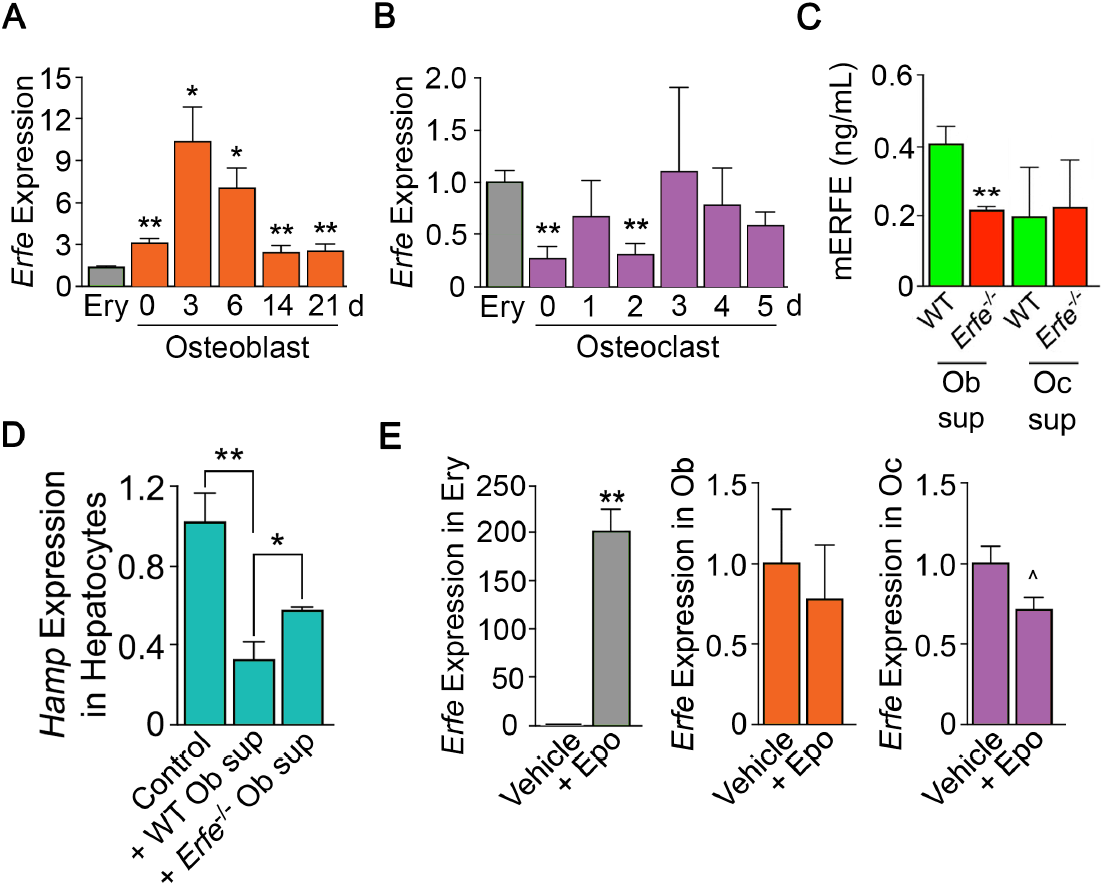
ERFE is Expressed at Higher Levels in Osteoblasts Than in Erythroblasts. (**A**) Quantitative PCR showing high levels of *Erfe* expression in osteoblasts from wild type mice cultured under differentiating conditions. Notably, at 3 days of culture, there was a 10-fold greater expression in osteoblasts relative to bone–marrow–derived wild type cultured erythroblasts. (**B**) Quantitative PCR showing similar Erfe expression in osteoclasts at 3-5 days of culture relative to bone–marrow–derived erythroblasts from wild type mice cultured under differentiating conditions. (**C**) Increased supernatant murine ERFE (mERFE) concentration in day 3 osteoblast cultures and no difference in day 5 osteoclast cultures from wild type relative to *Erfe*^-/-^ mice (detection limit of 0.2 ng/ml mERFE). (**D**) Hepcidin (Hamp) expression is suppressed in primary wild type hepatocytes in response to conditioned media from wild type relative to *Erfe*^-/-^ osteoblast cultures (day 6), confirming functionality of osteoblast-derived ERFE. Control hepatocytes were exposed to osteoblast culture media. (**E**) Unlike in erythroblasts, *Erfe* expression in cultured wild type osteoblasts and osteoclasts does not respond to erythropoietin (Epo). Statistics: Mean ± SEM; unpaired two–tailed Student’s *t*-test; *P<0.05, **P<0.01, ^0.05<P<0.1; *N*=3 wells/group.

To determine whether osteoblast–derived ERFE was functional, we established a bioassay based on the known inhibitory action of ERFE on hepcidin (*Hamp*) expression. For this, wild type hepatocytes were exposed to supernatants from differentiating wild type or *Erfe*^-/-^ osteoblasts. *Hamp* expression was suppressed with *Erfe*^-/-^ osteoblast supernatants, but importantly, this suppression was significantly greater with wild type supernatants (Fig. 2D). No *Hamp* suppression was evident with wild type or *Erfe*^-/-^ osteoclast supernatants. This latter suggests that osteoblast– but not osteoclast–derived ERFE is functional. However, as *Erfe*^-/-^ supernatants also suppressed *Hamp* expression, other yet unknown osteoblast–derived factors likely function in hepcidin regulation. Finally, unlike in erythroblasts, *Erfe* expression in mature osteoblasts or osteoclasts was not responsive to Epo (Fig. 2E).

Given that osteoblasts secrete ERFE that is known to inhibit hepcidin^21^ by sequestering BMPs^23,24,45^ that are skeletal anabolics^25^, we measured serum BMP2 concentration to find elevated BMP2 levels in *Erfe*^-/-^ relative to wild type mice (Fig. 3A). Given the specific importance of BMP2 in bone remodeling^26^, these results are consistent with the previously demonstrated sequestration of BMP2, along with BMP6, by ERFE^24^—namely, loss of ERFE led to decreased BMP sequestration. We thus hypothesized that ERFE functions in modulating bone formation by sequestering BMPs and tested whether the loss of ERFE facilitates BMP2–mediated signaling in the osteoblast *in vitro*. We found that the concentration of BMP2 was higher in supernatants from cultured *Erfe*^-/-^ osteoblasts compared with wild type cultures (Fig. 3B). Consistent with this difference, BMP2–activated signaling pathways, namely phosphorylated Smad1/5/8 and ERK1/2, but not phosphorylated p38, were enhanced in *Erfe*^-/-^ compared with wild type osteoblasts (Fig. 3C).

**Figure 3:**
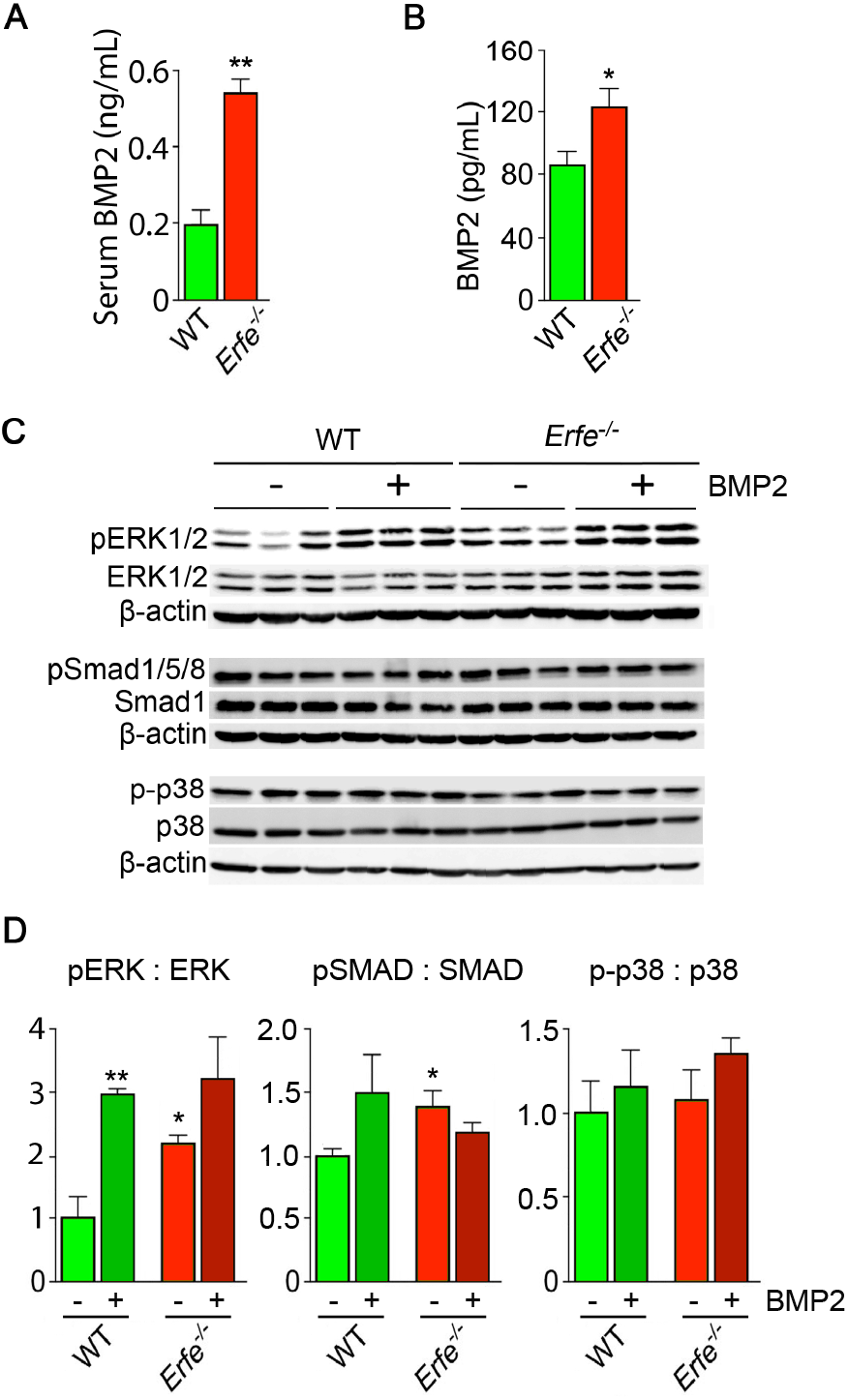
ERFE Function on Bone Involves BMP-2 Sequestration. (**A**) BMP2 ELISA demonstrates elevated BMP2 concentration in serum samples from *Erfe*^-/-^ relative to WT mice (*N*=4/group). In the 3–day cultures, there was an increase in BMP2 concentration in culture supernatants from *Erfe*^-/-^ relative to WT osteoblasts (*N*=6/group) (**B**). Similarly, signaling via the known BMP receptor pathways, namely ERK1/2 and Smad1/5/8, without changes in p38/MAPK; pSmad1/5/8 and pERK1/2 signaling was not further induced by BMP2 (50 ng/ml) in *Erfe*^-/-^ relative to WT osteoblasts. Western blots (**C**) with quantitation (**D**) shown. Statistics: Mean ± SEM; unpaired two–tailed Student’s *t*-test; *P<0.05, **P<0.01.

To further understand how ERFE impacts BMP2-mediated signaling, we evaluated the effect of BMP2 on wild type and *Erfe*^-/-^ osteoblasts *in vitro*. Treatment with BMP2 (50 ng/ml) in osteoblast cultures showed that pSmad1/5/8 and pERK signaling was not further induced in *Erfe*^-/-^ relative to wild type osteoblasts (Fig. 3C and 3D). In all, the data establish that increased BMP2 in *Erfe*^-/-^ mice leads to maximal induction of BMP signaling that remains unaffected by the further addition of BMP2. This supports the hypothesis that ERFE functions in bone by sequestering BMP2, thus, attenuating downstream signaling.

We studied whether the stimulation of bone formation in *Erfe*^-/-^ mice was due to a cell–autonomous action of ERFE on osteoblasts. For this, we compared the ability of wild type and *Erfe*^-/-^ bone marrow stromal cells *ex vivo* to differentiate into mature mineralizing colony forming units-osteoblastoid (Cfu-ob). Stromal cells from 5–month–old *Erfe*^-/-^ mice showed enhanced von Kossa staining of mineralizing Cfu-ob colonies (Fig. 4A). This mineralizing phenotype was associated with enhanced expression of the osteoblast transcription factors *Runx2* and *Sp7*, and downstream genes *Sost* and *Tnfsfll*^46,47^, increased supernatant RANKL levels, and suppressed expression of *Opg* (Figs. 4B, 4C). Enhanced RANKL profoundly increases osteoclastogenesis, as noted below, while sclerostin, encoded by the *Sost* gene, reduces the production of OPG, hence further increasing osteoclast formation.

**Figure 4:**
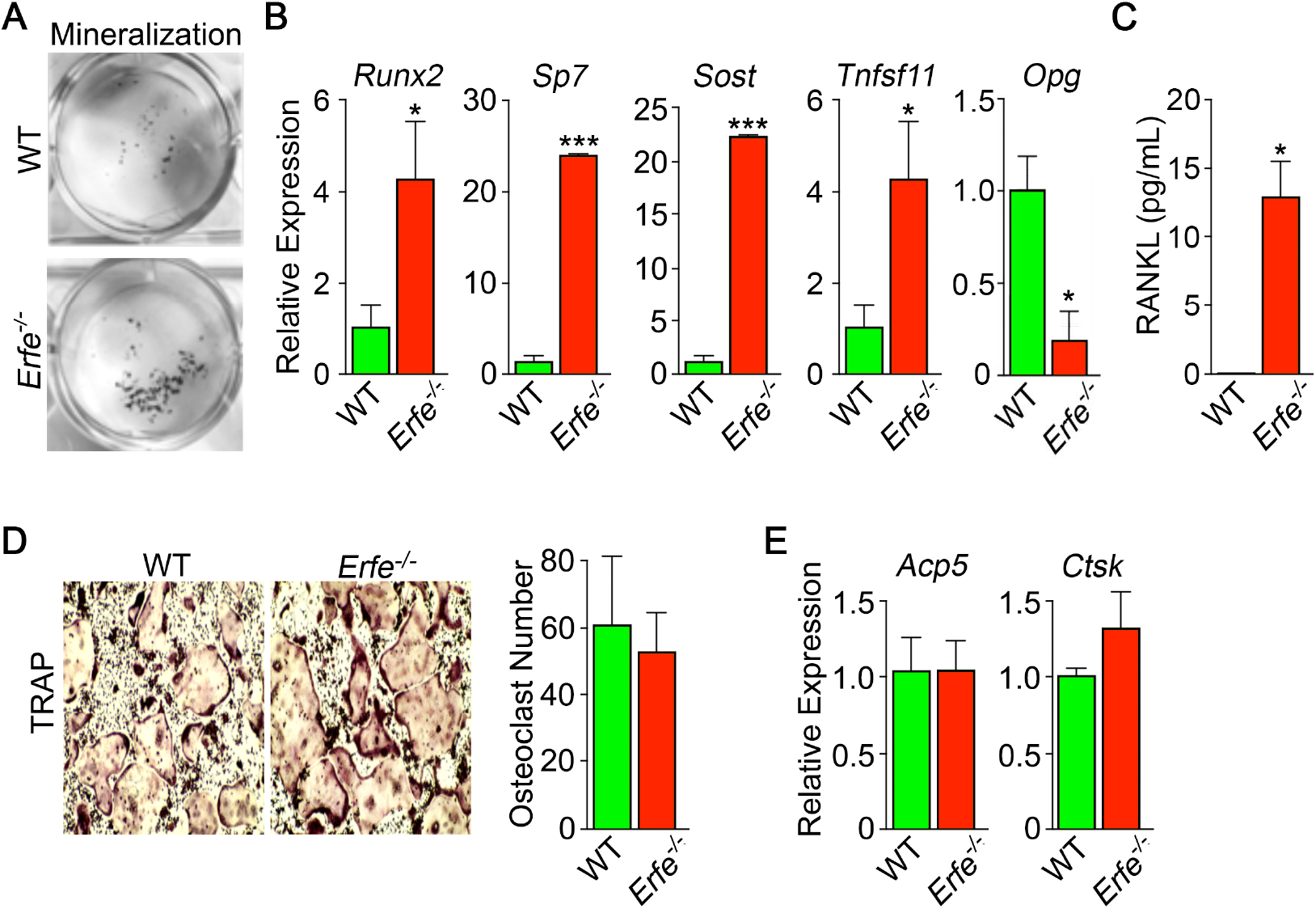
Mechanism of Action of ERFE on Bone Involves Interplay Between Osteoblastic RANKL and Sclerostin. Osteoblasts from 5–month–old wild type and *Erfe* mouse bone marrow cultured under differentiating conditions for 3 or 21 days. Loss of ERFE resulted in accelerated mineralization, noted by an increase in Von Kossa–stained nodules (**A**). Consistent with the cellular phenotype is the upregulation in *Erfe*^-/-^ osteoblasts of *Runx2, Sp7*, Sost and *Tnfsf11* expression and suppression of *Opg* expression (quantitative PCR on 21-day cultures) as well as (**B**) increased secreted RANKL (ELISA on 3–day cultures). (**C**) *In vitro* osteoclastogenesis assays show that ERFE loss does not alter osteoclast number, as measured by TRAP staining (**D**), or the expression of osteoclast genes, namely *Acp5* or *Ctsk* (**E**). Statistics: Mean ± SEM; unpaired two-tailed Student’s *t*–test; *P<0.05, **P<0.01; wells/group – 3 for A–C.

Given that *Erfe* is expressed in osteoclasts, and that *Erfe*^-/-^ mice display a pro-resorptive phenotype, we questioned whether ERFE directly affected the osteoclast, or whether the action resulted *via* a primary osteoblastic effect. *Erfe*^-/-^ bone marrow cell cultures derived from 5–month–old mice showed no difference in TRAP–positive osteoclast number compared to wild type cultures (Fig. 4D). Consistent with this, the program of osteoclast gene expression remained unchanged in these 5–day cultures (Fig. 4E). The data collectively suggest that the absence of ERFE results in the de-sequestration of BMP2, stimulates the osteoblast to upregulate RANKL and sclerostin, and thus enhances osteoclastic bone resorption indirectly.

Finally, we explored whether ERFE mediates osteoprotection in *Hbb*^*th3*/+^ mice, a model of human NTDT given that *Erfe* is upregulated in *Hbb*^*th3*/+^ marrow erythroblasts^3,21,37,48,49^. We crossed *Hbb*^*th3*/+^ mice with *Erfe*^-/-^ mice to generate *Hbb*^*th3*/+^;*Erfe*^-/-^ compound mutants. Whole body and site–specific measurements at mainly–cortical sites, namely femur and tibia, and vertebral trabecular (L4-L6) bone showed striking reductions in BMD in 5–month–old *Hbb*^*th3*/+^;*Erfe*^-/-^ mutants compared with *Hbb*^*th3*/wt^ mice, most notably in cortical bone (Fig. 5A). The trabecular bone loss was consistent with reduced histomorphometrically–determined fractional bone volume (BV/TV) and trabecular thickness (Tb.Th) in the femoral epiphyses (Fig. 5B). There was a trend towards increases in MAR (Fig. 5C), but a significant increase in TRAP–positive N.Oc and Oc.S in *Hbb*^*th3*+^;*Erfe*^-/-^ bones compared with wild type controls (Fig. 5D)—changes expected to produce reduction in bone mass. These findings document ERFE–mediated skeletal protection in β–thalassemia.

**Figure 5:**
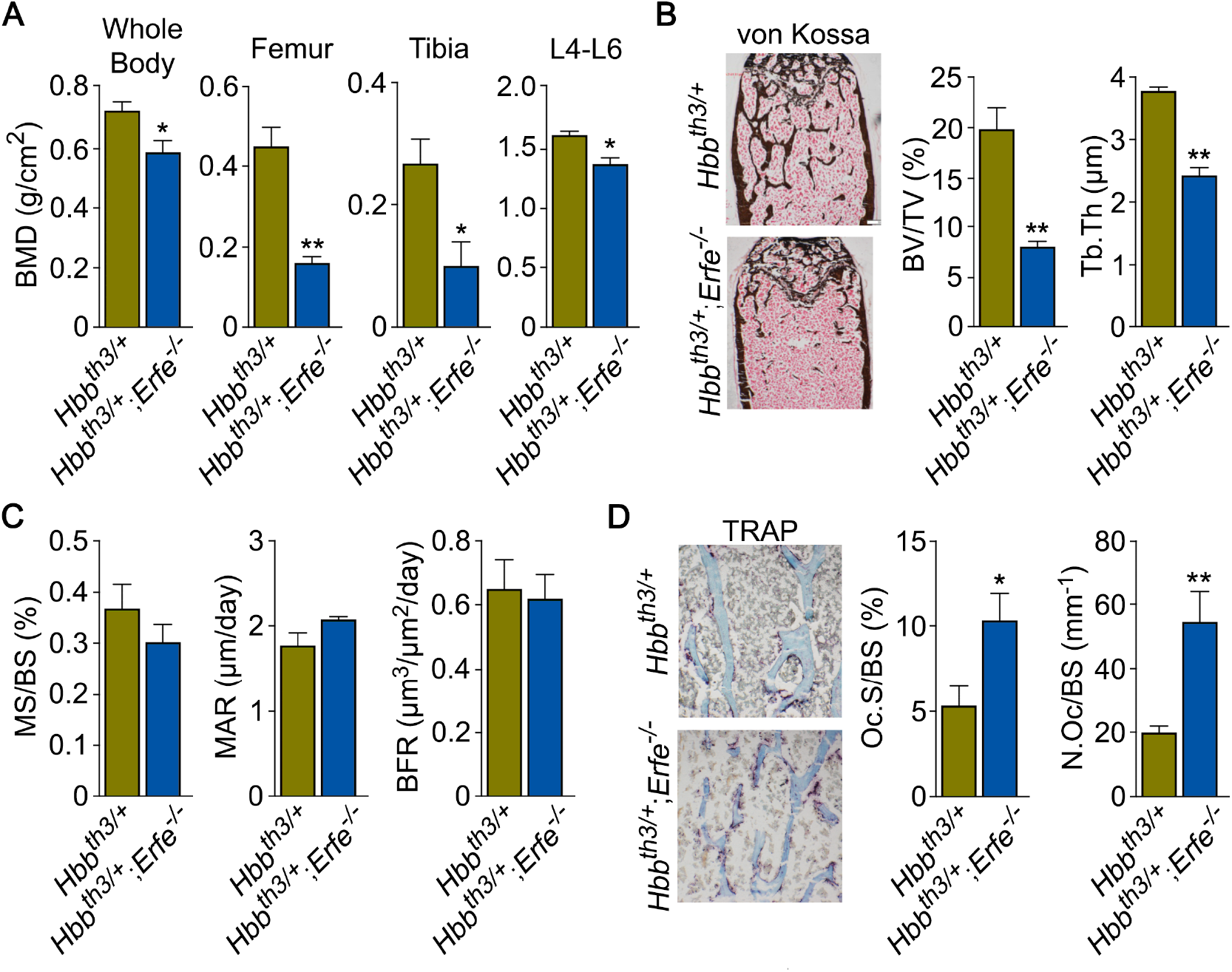
ERFE Loss in β-Thalassemia Mice causes Profound Bone Loss. (**A**) Bone mineral density (BMD) measured in whole body, femur, tibia, and lumbar spine (L4-L6) in 5–month–old β-thalassemia mice (*Hbb*^*th3*/+^ mice) and compound *Hbb*^*th3*/+^;*Erfe*^-/-^ mutants. (**B**) Representative section of femoral epiphyses stained with Von Kossa, and quantitative estimates of bone volume (BV/TV) and trabecular thickness (Tb.Th). (**C**) Dynamic histomorphometry following two i.p. injections of calcein (green) and xylenol orange (red) given at days −8 and −2, respectively. Shown are measured and derived parameters, namely mineralizing surface (MS), mineral apposition rate (MAR) and bone formation rate (BFR). (**D**) Representative image of TRAP (ACP5) staining of femoral epiphysis, also showing both osteoclast surface (Oc.S) and number (N.Oc), expressed as a function of bone surface (BS). Statistics: Mean ± SEM; unpaired two-tailed Student’s *t*-test; *P<0.05, **P<0.01; N=4–5 mice per group.

To confirm that decreased BMD in *Hbb*^*th3*/+^;*Erfe*^-/-^ mice did not result from further expanded erythropoiesis, we measured circulating red blood cells (RBCs) and reticulocytes, bone marrow erythroblasts, and spleen weight. Our results demonstrate a mildly decreased RBC count and hemoglobin, but no differences in spleen weight or bone marrow erythroblasts between 6–week–old *Hbb*^*th3*/+^ and *Hbb*^*th3*/+^;*Erfe*^-/-^ mice (Supplementary Fig. 2). This is consistent with what has been previously reported in *Hbb*^*th3*/+^;*Erfe*^-/-^ mice^43^.

## DISCUSSION

To date, the only known function of ERFE was on hepatocellular hepcidin expression exerted through the sequestration of BMPs^23,24^. Using genetically–modified mice and *in vitro* assays, we identify a new role for ERFE in skeletal protection. First, we show that *Erfe* expression is higher in osteoblasts comapred with erthyroblasts. Second, we find that ERFE is a potent down-regulator of BMP2–mediated signaling and RANK-L production by osteoblasts. Third, ERFE loss *in vivo* enhances bone formation, while also stimulating resorption by inducing the expression of osteoblastic *Tnfsf11* and *Sost*. The net effect of these opposing changes is bone loss in both young and old mice (Fig. 6). Fourth, although also produced by osteoclasts, ERFE displays no cell–autonomous actions on osteoclast function. Taken together, and consistent with prior inferences^27–32^, ERFE functions to protect the skeleton by negatively regulating BMP2 signaling in osteoblasts, with indirect inhibitory effects on osteoclastic bone resorption.

**Figure 6:**
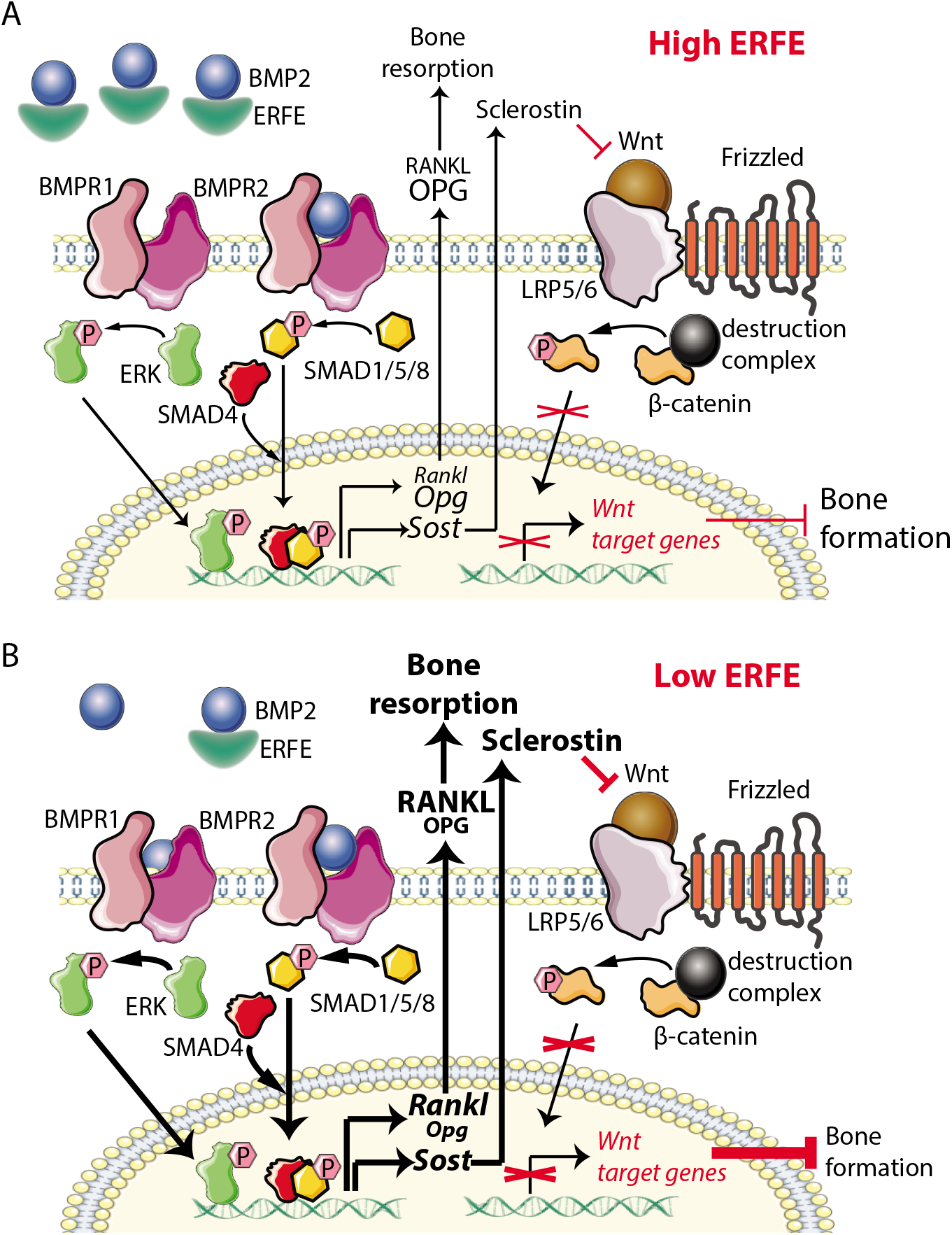
Putative Osteoprotective Function of ERFE in Health and in β–Thalassemia. In conditions of elevated ERFE (**A**), such as β–thalassemia, more BMP2 and BMP6 is sequestered, decreasing signaling through the BMP/Smad and ERK pathways. This would results in decreased *Sost* and *Rankl* expression to decrease osteoclastogenesis and bone resorption. In contrast, when ERFE is low (**B**), increased BMP2 leads to increased BMP/Smad and ERK signaling, increased *Sost* and *Rankl* expression and thereby, Sclerostin and RANKL release—this results in a greater suppression of Wnt signaling and increased osteoclastogenesis with consequent decrease in bone formation. Abbreviations: ERFE = erythroferrone; BMP = bone morphogenetic protein; BMPR = BMP receptor; SOST = sclerostin; RANLKL = receptor activator of nuclear factor kappa-B ligand; OPG = osteoprotegrin; LRP = lipoprotein receptor–related protein; Wnt = wingless-type MMTV integration site family.

We show that supernatants from wild type osteoblast cultures suppress *Hamp* expression, suggesting that BMP sequestration is likely a common mechanism that underpins both the hepatocellular and skeletal actions of ERFE. Recombinant ERFE binds certain bone–active BMPs, namely BMP2, BMP6, and the BMP2/6 heterodimer^23,24^, of which BMP2 is most relevant to adult bone formation^26^. We posit that ERFE is a negative regulator of osteoblastic bone formation, and that the absence of ERFE increases bioavailable BMP2 to promote osteoblast differentiation. Our results indeed demonstrate that BMP2 levels are elevated in supernatants from cultured *Erfe*^-/-^ osteoblasts, with enhanced downstream signals, notably phosphorylated Smad1/5/8 and ERK1/2. In all, the findings provide strong support for BMP sequestration as the mechanism of action of ERFE on bone.

We have also used the β–thalassemia mouse, *Hbb*^*th3*/+^, as a relevant disease model to study a role for ERFE in β–thalassemia, a condition with known elevations in ERFE. Chronic erythroid expansion in β–thalassemia is associated with a thinning of cortical bone resulting in bone loss^2,3^. It is therefore surprising that patients with the more severe forms of β–thalassemia, namely TDT, in whom RBC transfusions lead to suppression of endogenous erythropoiesis, exhibit significantly greater decrements in BMD than NTDT patients^49^. We have previously shown that ERFE is suppressed post-transfusion in TDT patients, and is significantly higher in NTDT patients^50^. Our finding of a marked reduction of bone mass in *Hbb*^*th3*/+^ mice with genetically–deleted *Erfe* (or *Hbb*^*th3*/+^;*Erfe*^-/-^ mice) compared with *Hbb*^*th3*/+^ mice provides strong evidence for a protective function of ERFE in preventing further worsening of the bone loss phenotype in β-thalassemia.

Taken together, our findings uncover ERFE as a novel regulator of bone mass *via* its modulation of BMP signaling in osteoblasts. In addition, because RBC transfusion suppresses erythropoiesis and thus decreases ERFE in both mice^48^ and patients^50^ with β-thalassemia, a relative decrement of ERFE may explain the more severe bone disease in TDT than in NTDT patients. As a consequence, our findings identify ERFE as a promising new therapeutic target for hematologic diseases associated with bone loss, such as β–thalassemia.

## ACKNOWLEDGEMENTS

We sincerely appreciate Tomas Ganz and Chun-Ling (“Grace”) Jung (UCLA), as well as Martina Rauner (University of Dresden) for many stimulating and helpful discussions and Ronald Hoffman (Icahn School of Medicine at Mount Sinai) for continued mentoring and advocacy. Y.Z.G. acknowledges the support of the National Institute of Diabetes and Digestive and Kidney Diseases (NIDDK) (R01 DK107670 to Y.Z.G. and DK095112 to R.F., S.R., and Y.Z.G.). M.Z. acknowledges the support of the National Institute on Aging (U19 AG60917) and NIDDK (R01 DK113627). S.R. acknowledges the support of NIDDK (R01 DK090554, R01 DK095112) and Commonwealth Universal Research Enhancement (C.U.R.E.) Program Pennsylvania.

## AUTHORSHIP CONTRIBUTIONS

Y.Z.G. conceptualized the study; M.C-M., S.G., M.R.M., A.G., M.P., C.C., V.S., M.F., C.C., S-M.K., D.L. performed and analyzed experiments; V.O., H.H., E.N., R.F., S.R., M.Z. and Y.Z.G. provided logistical support for these studies. T.Y., M.Z. and Y.Z.G. interpreted the data and wrote the manuscript. All authors participated in editing the manuscript.

